# Metformin-induced longevity is associated with retrotransposon dynamics in yeast chronological aging

**DOI:** 10.1101/2025.09.07.674784

**Authors:** Jimena Meneses-Plascencia, Ericka Moreno-Méndez, Diana Ascencio, Cesia Pérez-Aguilar, Michelle Munguía-Figueroa, Cei Abreu-Goodger, Soledad Funes, Alexander DeLuna

## Abstract

The widely used antidiabetic drug metformin extends lifespan across diverse model organisms, from yeast to primates. However, the cellular mechanisms underlying its anti-aging effects remain only partially understood. Here, we combined large-scale genetic screening and high-resolution lifespan phenotyping with transcriptomic and proteomic analyses to provide a systems view of metformin’s impact on the chronological lifespan of *Saccharomyces cerevisiae*. Unexpectedly, we uncovered pronounced gene-drug interactions between metformin and chromatin-modifying factors. Specifically, deletions of Set3C histone deacetylation complex subunits phenocopied the longevity effect of metformin, with no additive benefit when combined, suggesting convergence on shared pathways. Transcriptome profiling further revealed that metformin reprogrammed stationary-phase gene expression, with Ty1-copia retrotransposons emerging as a consistently induced signature, thereby suggesting a possible mechanism for the observed interactions with Set3C regulation. Paradoxically, TYA Gag-like protein levels and retrotransposition frequency were modestly reduced, indicating an uncoupling between transcriptional activation and retromobility. Proteome analysis revealed increased abundance of mitochondrial and stress-response proteins as primary outcomes of metformin exposure, both known modulators of Ty1 dynamics in yeast. Together, our findings position chromatin regulation and retrotransposon expression as integral components of metformin’s pro-longevity mechanisms, expanding its influence beyond signaling, metabolism, and stress response.

**Highlights:**

- Large-scale genetic screening reveals that deletions of Set3C histone deacetylase phenocopy metformin-induced longevity.
- Metformin consistently induces Ty1 retrotransposon transcription but reduces Gag protein abundance and retromobility.
- Proteomic changes highlight mitochondrial and stress-response proteins as primary outcomes of metformin exposure.
- Retrotransposon dynamics emerge as a key component associated with metformin-induced longevity.

## 1. Introduction

Metformin consistently extends lifespan in yeasts, nematodes, and mice (Anisimov et al., 2008; Borklu-Yucel et al., 2015; Onken and Driscoll, 2010; Seylan and Tarhan, 2023; Stynen et al., 2018). It also delays aging phenotypes and improves cognitive function in non-human primates (Yang et al., 2024). The geroprotective potential of metformin is further supported by evidence showing that it attenuates several hallmarks of aging, including impaired nutrient signaling, mitochondrial dysfunction, telomere attrition, cellular senescence, while also promoting autophagy (Barzilai et al., 2016; Kulkarni et al., 2020). Retrospective clinical studies have reported reduced mortality in diabetic patients treated with metformin (Bannister et al., 2014; Campbell et al., 2017). However, more recent analyses suggest that these associations may be affected by biases or confounding factors, contributing to an ongoing debate over the extent of metformin’s effects on human longevity (Stevenson-Hoare et al., 2023).

Despite its broad clinical use and pro-longevity potential, our understanding of the molecular mechanisms by which metformin modulates aging remains incomplete. Metformin is widely regarded as an activator of AMPK signaling, which has been linked to lifespan extension (Stancu, 2015; Zhou et al., 2001). It is also well established that metformin accumulates in mitochondria, where it acts as a mild inhibitor of complex I, altering the AMP/ATP ratio and triggering downstream effects including AMPK activation and TOR inhibition (Foretz et al., 2023; Pernicova and Korbonits, 2014; Rena et al., 2017). However, other mechanisms independent of AMPK have been described, including a lysosome-mediated pathway for AMPK and TOR regulation in *Caenorhabditis elegans* (Ma et al., 2022; Zhang et al., 2017), along with downstream outcomes such as improved nutrient sensing, mitochondrial biogenesis, SIRT1 activation, and increased autophagy (Kulkarni et al., 2020). In parallel, AMPK-independent effects of metformin also include reduced reactive oxygen species (ROS), anti-inflammatory signaling via NF-κB and Nrf2-Gpx7, and the activation of DNA damage response pathways such as ATM kinase (Kulkarni et al., 2020). The budding yeast *Saccharomyces cerevisiae* has emerged as a key model for studying the biology of aging (Kaeberlein, 2010), helping identify molecular factors that modulate aging and longevity in both this unicellular organism and more complex eukaryotes (Fabrizio and Longo, 2003; Janssens and Veenhoff, 2016). It also represents a powerful platform to characterize the mechanisms associated with potential anti-aging compounds (Zimmermann et al., 2018). However, few studies have systematically explored how metformin extends lifespan in yeast aging models. Transcriptome analyses in *S. cerevisiae* show that both glucose limitation and metformin enhance chronological lifespan (CLS), likely via improved mitochondrial function and mitochondria-to-nucleus signaling (Börklü, 2023; Borklu-Yucel et al., 2015). CLS extension by metformin in yeast is also linked to reduced advanced glycation end products and downregulation of mitochondrial respiration proteins (Kazi et al., 2017). Moreover, large-scale homomer dynamics screens suggest that metformin triggers a cellular response resembling iron deficiency (Stynen et al., 2018). In the fission *Schizosaccharomyces pombe*, metformin increases glucose consumption and ATP production while lowering ROS, indicating enhanced metabolism and oxidative stress resistance (Seylan and Tarhan, 2023). Still, an integrated mechanistic view remains elusive, and key molecular players may yet be unidentified, given the widely recognized pleiotropic nature of metformin’s cellular effects.

In this study, we investigated the mechanisms underlying metformin-induced chronological longevity of yeast cells by combining large-scale genetic screening with transcriptomic and proteomic analyses. We found that metformin extends CLS through mechanisms independent of pH buffering, involving stress responses, mitochondrial function, and an unexpected involvement of the SET3C histone deacetylase complex, which had not been previously linked to metformin’s effects. Notably, we identified a novel connection between metformin treatment and Ty1 retrotransposon regulation, whereby metformin increased Ty1 expression without promoting its mobility, suggesting that retrotransposon dynamics contribute to its anti-aging effects.

## 2. Materials and Methods

### 2.1. Strains and media

Large-scale competitive-aging screening was carried out with a set of 1438 strains deleted for genes with at least one ortholog in human. Orthologs were defined with the BioMart tool from Ensembl (Kinsella et al., 2011). A collection of haploid strains tagged with mCherry red fluorescent protein (RFP) was generated by crossing viable BY4741 deletion strains (*MAT*a *xxx*Δ::*kanMX4 his3*Δ*1 ura3*Δ*0 leu2*Δ*0 met15Δ0*) with the Y8205-RFP SGA-starter strain (*MAT*α *PDC1-RFP-CaUR*A3 *his3*Δ*1 ura3Δ0 can1*Δ*::STE2pr-SpHIS5 lyp1*Δ). The resulting prototrophic haploid progeny had the following genotype: *MAT*a *xxx*Δ*::kanMX4 PDC1-RFP-CaURA3MX4 can1*Δ*::STE2pr-SpHIS5 lyp1*Δ *LEU2 his3*Δ*1 ura3*Δ*0*. WT reference strains had insertions at the neutral *his3*Δ locus, tagged with Cerulean (CFP) or mCherry (RFP). The *RFP*-tagged *his3*Δ reference was also used as the WT for transcriptomic profiling by RNA-seq.

For large-scale flow-cytometry proteomics, a subset of 2600 strains from the GFP fusion collection (Huh et al., 2003) was used; selection included all strains with detectable flow-cytometry GFP signal (Newman et al., 2006). Selected GFP strains were crossed with the strain *Mat*α *ho*Δ::*BFP-G418 HIS3::HYG* resulting in an array of strains with genotype: *Matα XXX-GFP-HIS3Mx6 ho*Δ::BFP-G418 *can1Δ::CaUra3 lyp1Δ::STE3pr*_*LEU2 his3*Δ::*HYG*.

All chronological lifespan assays were carried out in synthetic complete medium (SC) medium buffered to pH 4 (citrate-phosphate buffer), unless otherwise noted. SC medium contained 2% glucose, 0.2% complete amino acid mix, 0.67% yeast nitrogen base (YNB) without amino acids, 30 mM citric acid, and 18.5 mM sodium citrate. Where indicated, metformin (Sigma D150959) was added to the medium to final concentrations ranging from 3 to 200 mM. Outgrowth medium was low-fluorescence YNB (Sheff and Thorn, 2004).

### 2.2. High-throughput CLS screening by competitive-aging

Mutant-RFP arrays and WT-CFP were grown separately until saturation and then mixed in a 2:1 ratio. Mixed cultures were replicated by pinning into a 96 semi-deep-well with 750 µL SC medium (pH 4), with or without 50 mM of metformin. Cultures were incubated without shaking at 30 °C and were sampled during stationary phase for nine consecutive days, starting at 60 h after inoculation. Ten microliter aliquots were used to inoculate 140 µL of fresh YNB low fluorescence medium employing a 96-channel pipetting arm. Outgrowth kinetics were monitored every 1.5 h for 15 h by recording absorbance at 600 nm (OD600*nm*) along with raw fluorescence of *RFP* (Ex 587/5 nm, Em 610/5 nm) and *CFP* (Ex 433/5 nm, Em 475/5 nm) using a Tecan Infinite M1000 reader integrated to a cell-assay robotic platform (Tecan Freedom EVO200). Optimal gain was calculated before the start of each batch experiment.

### 2.3. Data normalization and analysis

The survival coefficients (*S*) of each sample relative to the WT reference was estimated by fitting the data to a multiple regression model, as reported (Avelar-Rivas et al., 2020). MATLAB scripts are available at https://abrahamavelar.github.io/LinearModelCLS/. Raw survival coefficients were normalized for batch effects by subtracting the median *S* of each plate. The final CLS normalized dataset is provided in **Table S1**. CLS screening also included a total of 227 reference samples, in which an RFP-tagged WT was aged in competition with a CFP-tagged WT. The distribution of these reference samples was used to compute a Z-score and associated *p*-value for each mutant tested (normally distributed *x*^2^ test, *p*=0.16 and *p*=0.29 for ‒MTF and +MTF, respectively). Short- and long-lived phenotypes in each condition were defined after correcting for multiple hypotheses testing (*FDR, q*<0.05).

Lifespan extension by metformin (*LE_MTF_)* was defined as:

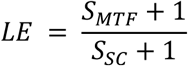

Likewise, diminished or enhanced lifespan extension was determined by assigning each mutant a *Z*-score from the *LE* distribution of the 227 WT reference replicates (normally distributed, *p*= 0.19, *x*^2^ test) and corrected for multiple hypothesis (*q*<0.05, Benjamini–Hochberg FDR).

### 2.4. Functional enrichment analysis

To identify gene sets with atypical *LE* median values, all genes tested were grouped by their annotated GO terms; only those groups with at least 5 genes and no more than 50 genes measured were considered. Annotation files were downloaded from the Yeast Genome Database (2023). The distribution of *LE* values within each group was compared to the distribution of all deletion strains tested. Gene sets associated with longevity by metformin were defined as those with a median *LE* value different to the entire data set (Wilcoxon rank sum test, p<0.05). Significant gene sets and the genes within each set are listed in **Table S2**.

### 2.5. Validation of CLS phenotypes by live/dead staining

Selected strains were pre-cultured for 36 h in 96-well plates (Corning 3585). Saturated cultures were pinned into 96 semi-deep-well plates containing 750 μL SC medium (pH 4), with or without metformin. Cultures were incubated without shaking at 30 °C. After 48 h, 6 μL samples were taken using a Tecan Freedom EVO200 robotic arm and transferred to 96-well plates with 50 μL of staining solution (3.34 μM propidium iodide and 20 μM Syto9; LIVE/DEAD™ FungaLight™ Yeast Viability Kit). Live cells were quantified by flow cytometry (LSRFortessa™, BD), using Syto9-positive events. Propidium iodide was excited at 561 nm (586/15 BP filter), and Syto9 at 488 nm (525/50 BP, 505LP filters). Survival curves were fitted with an exponential decay model to estimate half-life, and survival integrals were calculated as the area under the curve for each independent experiment. Statistical differences of 50 mM treatment compared to the ‒MTF control were assessed by unpaired two-tailed Student’s t-test (*p* < 0.05).

### 2.6. Proteomic response to metformin measured by analytical flow cytometry

GFP-tagged strains were pre-cultured for 36 h in 96-well plates (Corning 3585). Using a Tecan Freedom EVO200, 10 μL of each saturated culture was transferred into 96 semi-deep-well plates containing 750 μL SC medium (pH 4), with or without metformin. Cultures were incubated at 30 °C without shaking for 24 h, 48 h, or five days, as indicated. At each time point, 10 μL samples were mixed with 50 μL Tris-EDTA buffer in 96-well plates and analyzed by flow cytometry (LSRFortessa™, BD). GFP was excited with a 488 nm laser, and emission was detected using a 525/50 band-pass filter. Doublets were excluded using side scatter (SSC), and median GFP fluorescence was calculated from singlet events. Log2 fold changes were estimated as the protein-abundance change ratio under each treatment. Functional analysis was performed as described previously for *LE* values, by comparing the distribution of Log2 fold changes of each gene set to that of the complete set of 2616 GFP-fusion strains sampled.

### 2.7. RNAseq and data analysis

Pre-inoculums of the WT strain were established by culturing 5–6 individual colonies in 10 mL of SC medium (pH 4) at 30 °C with shaking (200 rpm) for 24 h. Cultures were then diluted to an OD₆₀₀ of 0.05 in 50 mL of SC medium (pH 4), with or without metformin, and incubated under the same conditions. Cells were collected at mid-exponential phase (OD₆₀₀ = 0.5) and late stationary phase (8 days), with 10⁷ cells/mL harvested per replicate. Total RNA was extracted using the RiboPure™ Yeast Kit (Invitrogen AM1926). Four replicates from the exponential phase and five from the stationary phase were used for directional, rRNA-depleted, 150 bp paired-end sequencing (Novogen).

Low-quality bases and adapter sequences were trimmed using FASTP. Trimmed reads were aligned to the reference genome using STAR and filtered by mapping quality with Samtools. Read counts were obtained using HTSeq. Data normalization was performed with the RUVseq package, and differential expression analysis was conducted using DESeq (FDR < 0.1, Benjamini-Hochberg) in RStudio. GO over-representation analysis was carried out using the enrichGO function from the clusterProfiler package. A Jaccard index and hierarchical clustering were applied to the resulting GO terms, and functional networks were constructed in Cytoscape (v3.10.2).

### 2.8. Retromobility assay

A Ty1his3-AI retrotransposon reporter strain was obtained by lithium-acetate transformation of the BY4741 strain with a centromeric plasmid construction that contains a Ty1 element with its native promoter (Hénault et al., 2020). The construction carried the YMRCTy1-3 Ty1 element (YMR045C). The TYA Gag and TYB Pol coding regions of this LTR retrotransposon were chosen because of their significant upregulation by 50mM metformin compared to the ‒MTF control (0.47 log2 fold change; *p*=0.002, Benjamini-Hochberg FDR).

Ty1 his3-AI mobility frequency was assayed as previously described (Nyswaner et al., 2008), with modifications. Starting cultures of the Ty1 reporter strain were initiated by inoculating 10 mL of SC medium (pH 4) with three isolated colonies and incubating at 30 °C with shaking (200 rpm) for 24 h. Saturated cultures were diluted 1:1000 into 25 mL of fresh SC medium (pH 4), with or without metformin, and incubated at 20 °C with shaking (100 rpm) for 75 h to allow retrotransposition. To estimate colony-forming units (CFUs), serial dilutions were prepared and 100 μL of the 10⁻⁵ dilution was plated on YPD agar. In parallel, 12 mL of each culture was centrifuged (2000 × g, 10 min, RT), washed twice, and resuspended in 2 mL of sterile water. Cells were plated on SC–His agar using a sterile glass spreader and incubated at 30 °C for 48 h (CFU) or 72 h (His⁺ colony counting). Retrotransposition frequency was calculated by dividing the number of His⁺ colonies by CFUs per μL.

## 3. Results

### 3.1. Metformin increases chronological longevity under pH-buffered conditions

Metformin has been shown to increase the chronological longevity of *S. cerevisiae* (Borklu-Yucel et al., 2015; Stynen et al., 2018). However, metformin also prevents pH acidification of stationary phase cultures in conventional SC medium (Avelar-Rivas et al., 2020). Since acetic acid-induced mortality is a major driver of chronological aging under non-buffered conditions (Burtner et al., 2009; Ludovico et al., 2001), the buffering effects of metformin may explain its impact on chronological survival. We thus set out to evaluate the effects of metformin on the CLS phenotype of cells aged under buffered-media conditions. To determine an optimal pH for CLS profiling, we first evaluated the effects of the drug on the growth kinetics of a WT strain grown under SC media buffered at a range of pH (**Figure S1**). We observed that high doses of metformin had little to no effect on cell growth at a pH range of 3 to 5, while cells grown at higher pH values showed increased sensitivity as a function of drug concentration. Based on this result, we reasoned that stationary-phase survival would be better assayed in media buffered at pH 4 in which cells grow at their normal rates.

To assess the chronological-aging effects of metformin under buffered conditions, we used live/dead staining coupled to flow cytometry to characterize the CLS of cells aged on SC medium buffered at pH 4 under increasing concentrations of the drug (**Figure 1A**). We observed that metformin led to increased chronological longevity in a dose-dependent manner. Specifically, the half-life of the WT increased from 25 days in the control to up to 41 days in the 75 mM treatment (39% CLS increase) (**Figure 1B**). We also observed that metformin was rendered deleterious at high doses, with a half-life reduction to 17 days in the 200 mM treatment (41% decrease). These results show that metformin can increase chronological longevity in buffered conditions, suggesting that part of the cellular mechanisms of longevity by metformin in yeast are independent of its buffering capacity.

**Figure 1.**
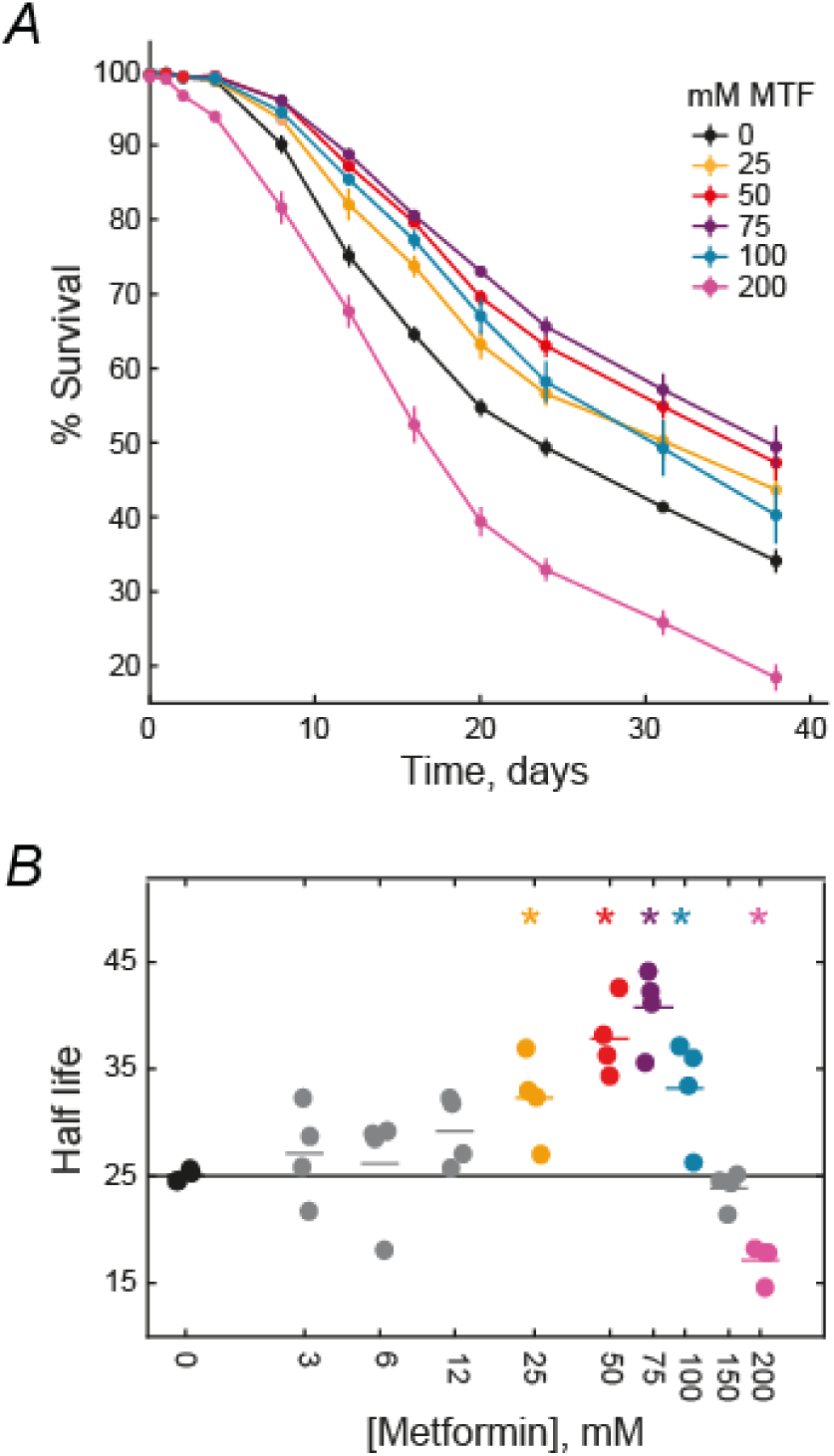
Metformin extends CLS in a dose-dependent manner. ***A*)** The WT strain was aged with increasing metformin concentrations in SC medium buffered at pH 4 and survival was tracked by live and death staining coupled with flow cytometry. The plot shows only those treatments that resulted in increased or decreased CLS. Error bars are the S.E.M. (n=4). ***B*)** Survival half-lives were estimated by fitting the survival data to an exponential decay model; asterisks indicate treatments that were statistically different to the control without metfrormin (*p*<0.05, unpaired two-tailed *t*-test).

### 3.2. Large-scale genetic profiling of aging in metformin

To systematically identify genetic factors underlying lifespan extension by metformin in budding yeast, we carried out a large-scale functional genomics aging assay. Specifically, we screened a set of 1438 knockout strains from the yeast deletion collection, which included most non-essential genes having at least one human ortholog (Kinsella et al., 2011). To this end, we used a sensitive competitive-aging assay developed in our laboratory, which tracks changes in stationary-phase viability of gene-deletion strains aged in coculture with a fluorescently tagged WT reference (**Figure 2A**) (Avelar-Rivas et al., 2020; Campos et al., 2018; Garay et al., 2014). With this approach, we obtained a relative survival coefficient for each knockout strain aged with 50 mM metformin (+MTF) or without the drug (–MTF) (**Figure 2B**). Survival coefficients were normalized to account for batch effects by subtracting the median raw value of each plate (**Figure S2**). Importantly, a subset of 84 mutant strains was assayed under both conditions in five independent experiments, showing high replicability in the survival coefficients measured (*r* =0.90–0.97; **Figure S3**). All large-scale genetic screening data is provided in the Supporting Information (**Table S1**).

**Figure 2.**
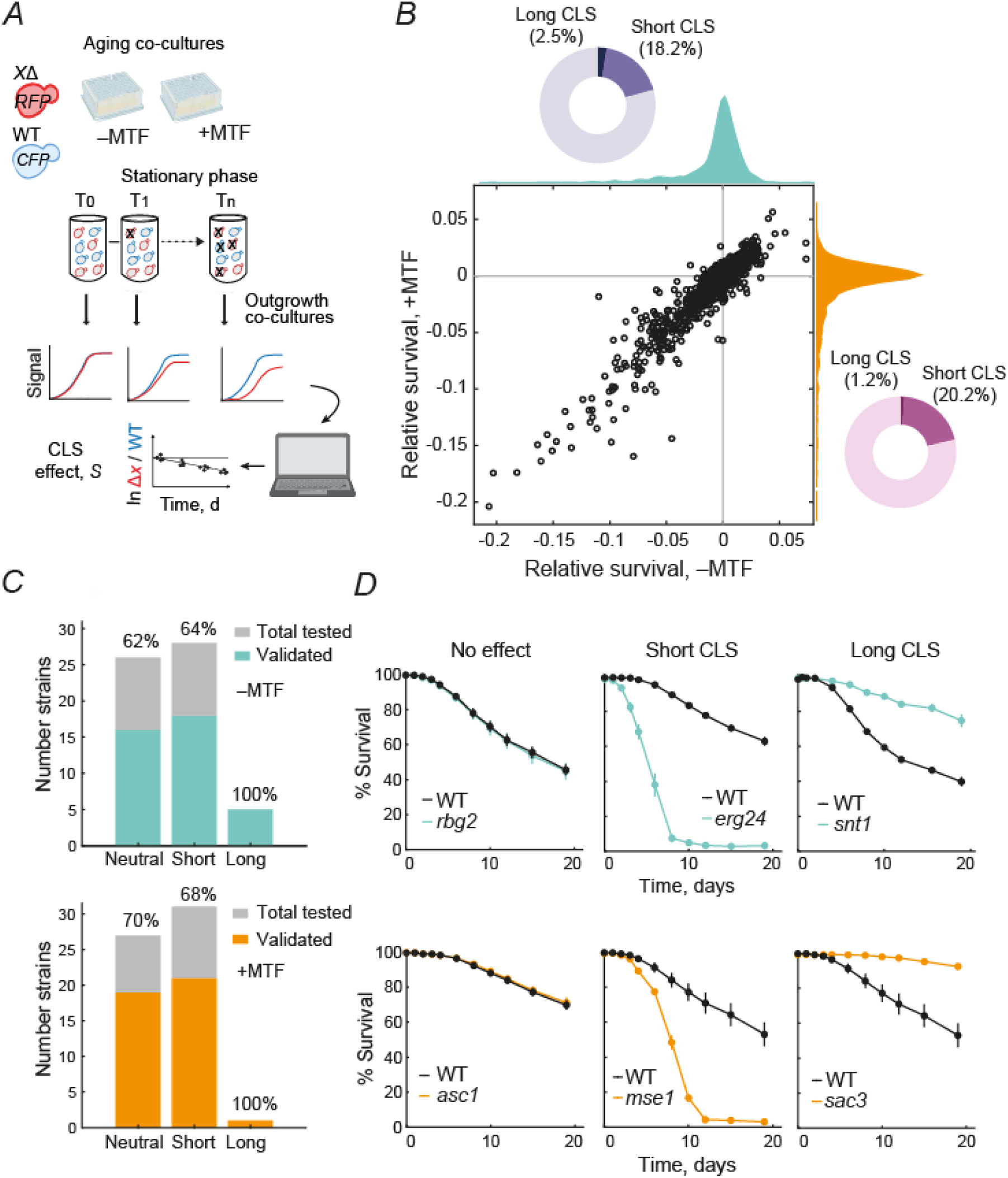
High-resolution genetic screening of lifespan extension by metformin. ***A)*** Schematic representation of the competitive method used to test the effect of metformin on *CLS*. Saturated cultures of *RFP*-labeled deletion strains and *CFP*-labeled *WT* strain are mixed and inoculated into SC medium without metformin (‒MTF) or 50 mM metformin (+MTF). Samples of the aging co-cultures are taken over the days and inoculated in a fresh medium. The outgrowth kinetics are monitored by recording the optical density and tracking the fluorescence signals. The collected data is fitted to a multiple linear regression model to estimate the relative survival coefficient (CLS effect, *S*) for each mutant tested. ***B***) The scatter plot displays the relative survival coefficients of 1438 knockout strains of human ortholog genes aged in a medium without metformin (‒MTF, horizontal axis) and supplemented with 50 mM MTF (+MTF, vertical axis). The orange and cyan histograms correspond to the distribution of the survival coefficients in ‒MTF and +MTF, respectively. Pie charts display the percentage of long and short CLS strains detected in each treatment. CLS phenotypes were identified through *Z*-score estimation, by comparing the mutant’s survival coefficient to the survival distribution of 227 wild-type reference strains. A multiple-testing correction was then applied to define hits at a 5% FDR (Benjamini–Hochberg). ***C*)** Percentage of validated strains by live and death assay shown by CLS phenotype in ‒MTF (cyan) and +MTF (orange). ***D*)** Representative survival curves obtained by live and dead assays of short, long and neutral strains compared to a wild-type strain (black line) aged in ‒MTF (cyan lines) or +MTF (orange lines). Error bars are the S.E.M. (*n*=5).

To score gene-deletion strains with increased or decreased CLS under each treatment, the distribution of the survival coefficients of all deletion strains was compared to a null distribution of 227 WT strains. In the control SC medium without the drug, 261 (18.2%) deletion strains showed significantly decreased lifespan, while 36 (2.5%) mutants showed increased lifespan compared to the WT (**Figure 2B**). The number of significant CLS effects in the +MTF treatment were 291 (20.2%) and 17 (1.18%) for short- and long-lived deletion strains, respectively (**Figure 2B**).

To validate results of our CLS genetic screens, we used live/dead staining to estimate the CLS phenotype of a subset of deletion strains assayed at smaller scale. This set included strains that were top hits under at least one of the two conditions, which also allowed testing for false negatives given that many of the CLS phenotypes were only observed under one condition. Our validation experiments showed that, without metformin, 16 out of 26 (62%) strains recapitulated their neutral CLS phenotype, while 18 / 28 (64%) and 5 / 5 (100%) recapitulated their short- and long-lived CLS phenotypes, respectively, when tested by live/dead staining (**Figure 2C**). Likewise, 19 / 27 (70%), 21 / 31 (68%), and 1 / 1 (100%) deletion strains showed consistent neutral, short-, and long-lived CLS phenotypes when retested in the presence of metformin. Selected validation experiments are shown in **Figure 2D**; all hits tested are provided in **Figure S4**. The rate of validation of hits in our large-scale screens is similar to that reported in previous CLS surveys (Campos et al., 2018; Matecic et al., 2010). These results indicate that our competitive aging approach yields an accurate estimation of CLS under different conditions, allowing us to further describe gene-drug interactions and their role in lifespan extension by metformin.

### 3.3. Altered chromatin organization phenocopies the longevity effect of metformin

To gain deeper insight into the genes and cellular processes underlying metformin’s lifespan-extending effect, we analyzed our large-scale genetic data to identify deletions that alter survival coefficients under metformin treatment relative to drug-free medium. To this end, we calculated a relative lifespan extension coefficient (*LE_MTF_*) for each deletion strains, which provides a metric to identify genes associated with diminished or enhanced relative lifespan extension upon treatment with metformin. Using a 5% FDR cutoff, we found that 45 deletion strains showed decreased *LE_MTF_*, whereas 24 lived significantly longer under metformin treatment (**Figure 3A**). Deletion of *TPK1*, which encodes a catalytic subunit of PKA, showed diminished lifespan extension, whereby its short-lifespan phenotype was strongly aggravated by metformin. This supports a requirement for intact PKA signaling in metformin-induced longevity, consistent with its archetypical mechanisms of action. Unexpectedly, other top hits with diminished lifespan extension included strains impaired in chromatin regulation. For example, deletions of *SNT1* or *HST1*, coding for subunits of the Set3C histone deacetylase complex, extended lifespan in drug-free medium but were rendered neutral or detrimental in the presence of metformin. These kind of alleviating gene-drug interactions suggest that the longevity effects of gene deletions and metformin treatment arise from common underlying mechanisms.

**Figure 3.**
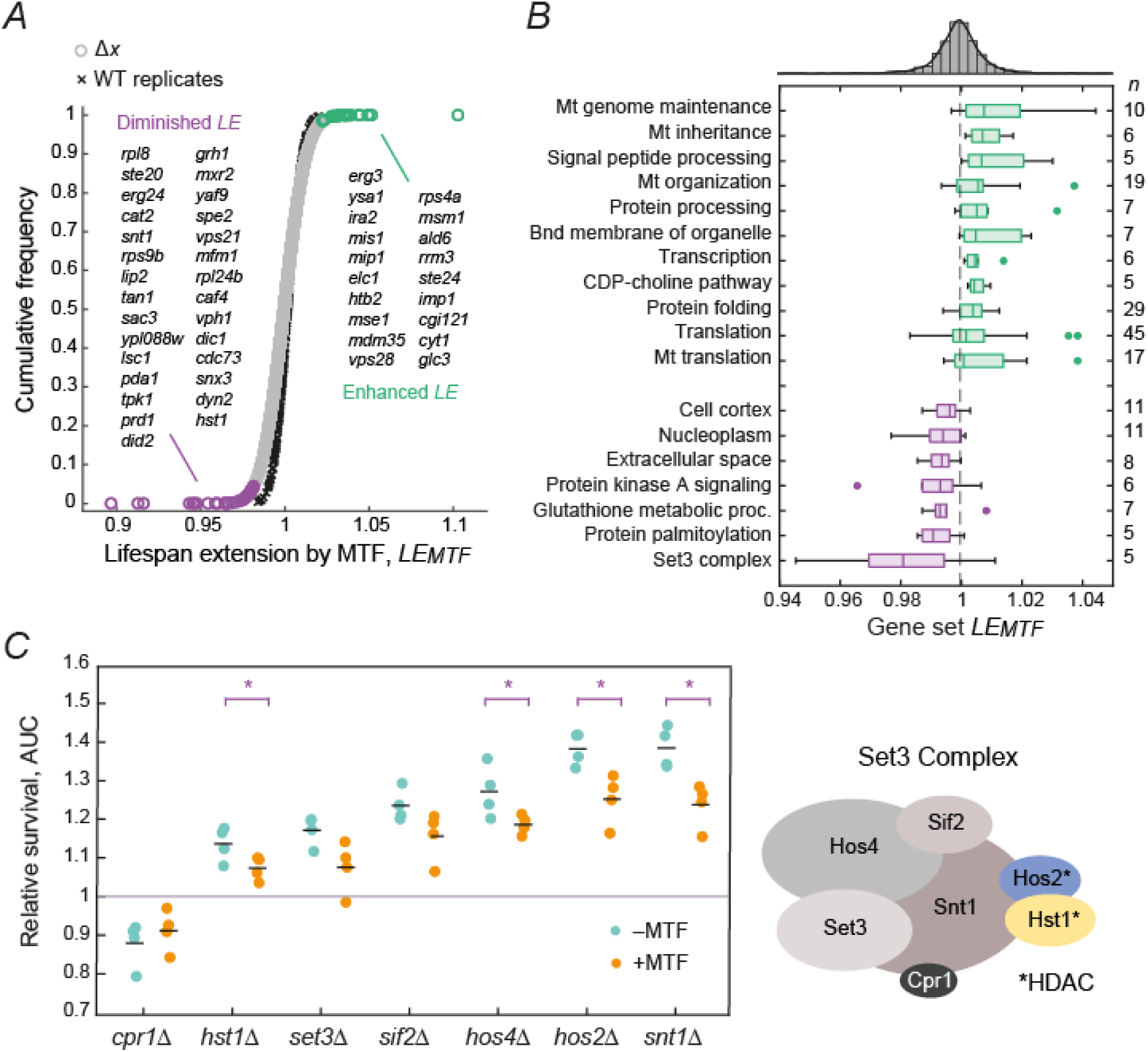
Genetic interactions underlying metformin-induced longevity. ***A*)** Lifespan extension by metformin, *LE_MTF_*, was calculated for each mutant (grey circles) and compared to the *LE_MTF_* distribution of WT strains (black crosses) to score statistical significance at 5% FDR (highlighted in purple and green for diminished and enhanced CLS extension, respectively). Top hits (1% FDR) are displayed in the plot. ***B*)** The boxplots show the *LE_MTF_* (horizontal axis) of genes annotated within the same GO term (vertical axis); *Mt* indicates mitochondrial-related categories. CLS specific gene groups exhibiting an increase in median lifespan are plotted in green, while gene groups with a reduced median *LE_MTF_* are shown in purple (Wilcoxon rank sum test). The dashed gray line is the median *LE_MTF_* of all strains tested (*n* = 1438), while the histogram on the top shows the data distribution of the entire dataset. ***C*)** Validation of hits for genes encoding subunits of Set3C. Replicates of single and double mutants were aged in individual cultures, and survival curves were obtained using live/dead staining of stationary-phase cultures at different days. The plot shows the area under the curve (AUC) for the mutants, expressed relative to the average AUC of the WT aged in ‒ MTF (cyan) or +MTF treatment (orange). Statistical differences between each treatment pair were assessed by unpaired two-tailed *t*-test (\**p*<0.05). Set3C schematic adapted from (Ryu, 2024); asterisks indicate protein subunits with annotated histone deacetylase activity.

To better resolve the cellular functions underlying metformin-induced longevity, we used our quantitative dataset to search for GO terms with significant skew in their distributions of *LE_MTF_* values. This quantitative analysis avoids reliance on arbitrary cutoffs and can reveal functional groups enriched in genes with milder effects that would be missed by standard significance thresholds. Using this strategy, we identified several processes consistently linked to metformin action across model organisms, including PKA signaling, mitochondrial function, and redox metabolism (**Figure 3B**; **Table S2**). In addition, cellular proteostasis, vacuole function, and the extracellular space were also hits showing significant deviation in their median *LE_MTF_* values. Noteworthy, impairment of Set3C chromatin modification proved to be the top hit of gene deletions consistently associated with diminished longevity by metformin.

We focused on the unexpected link of the Set3C histone deacetylation complex with the cellular response to metformin, which was the top functional hit enriched in negative gene-drug interactions. In *S. cerevisiae*, this complex includes 7 non-essential subunits; it is the yeast analog of the mammalian HDAC3/SMRT complex (Pijnappel et al., 2001). Using live/dead staining of stationary-phase cells aged in individual cultures, we re-evaluated the genetic interactions of all Set3C subunit deletion strains aged with and without metformin (**Figure 3C**). This validation experiment included single deletions of *SET3* and *HOS2*, which were not part of the original large-scale screen due to the absence of direct human orthologs. With the sole exception of Δ*cpr1*, all Set3 complex subunits mutants showed diminished lifespan extension, whereby the strong long-lived phenotypes of the deletion strains were partially suppressed in cells aged with metformin. Significant negative gene-drug interactions—defined as diminished lifespan extension in response to metformin—were observed for *HOS4*, the histone deacetylases *HST1* and *HOS2*, and *SNT1*, the scaffold component of Set3C. This pattern of diminished lifespan extension suggests that impaired Set3C chromatin regulation and metformin treatment act through overlapping pathways that promote chronological longevity. In this context, Set3C deletions phenocopy the longevity effect of metformin, and no additive effect is observed when both interventions are combined. Our large-scale genetic interaction analyses reveal consistent and reproducible gene-drug interactions between Set3C components and metformin.

### 3.4. Metformin leads to increased expression of Ty1 retrotransposons in aging cells

To shed light into the nature of the overlapping regulatory pathways linking the cellular responses to metformin and impaired Set3C chromatin, we carried out a global transcriptomic analysis. Specifically, we performed RNAseq analysis comparing populations of WT cells in mid exponential and late stationary phase with or without metformin. As expected, PCA analysis of replicated samples showed a strong signal clustering stationary-phase cells apart from exponential-phase cells, and a moderate clustering of metformin treated cells, but only in stationary phase (**Figure 4A**). Despite its impact on chronological lifespan, metformin had a relatively modest effect on the stationary-phase transcriptome, with significant expression changes detected in only 3.0% of genes. After performing a differential expression analysis for stationary phase cells, we found that metformin resulted in significant upregulation of 154 genes and downregulation of 25 genes (*p*<0.05, Benjamini-Hochberg FDR adjusted) **(Figure 4B**; **Table S3**).

**Figure 4.**
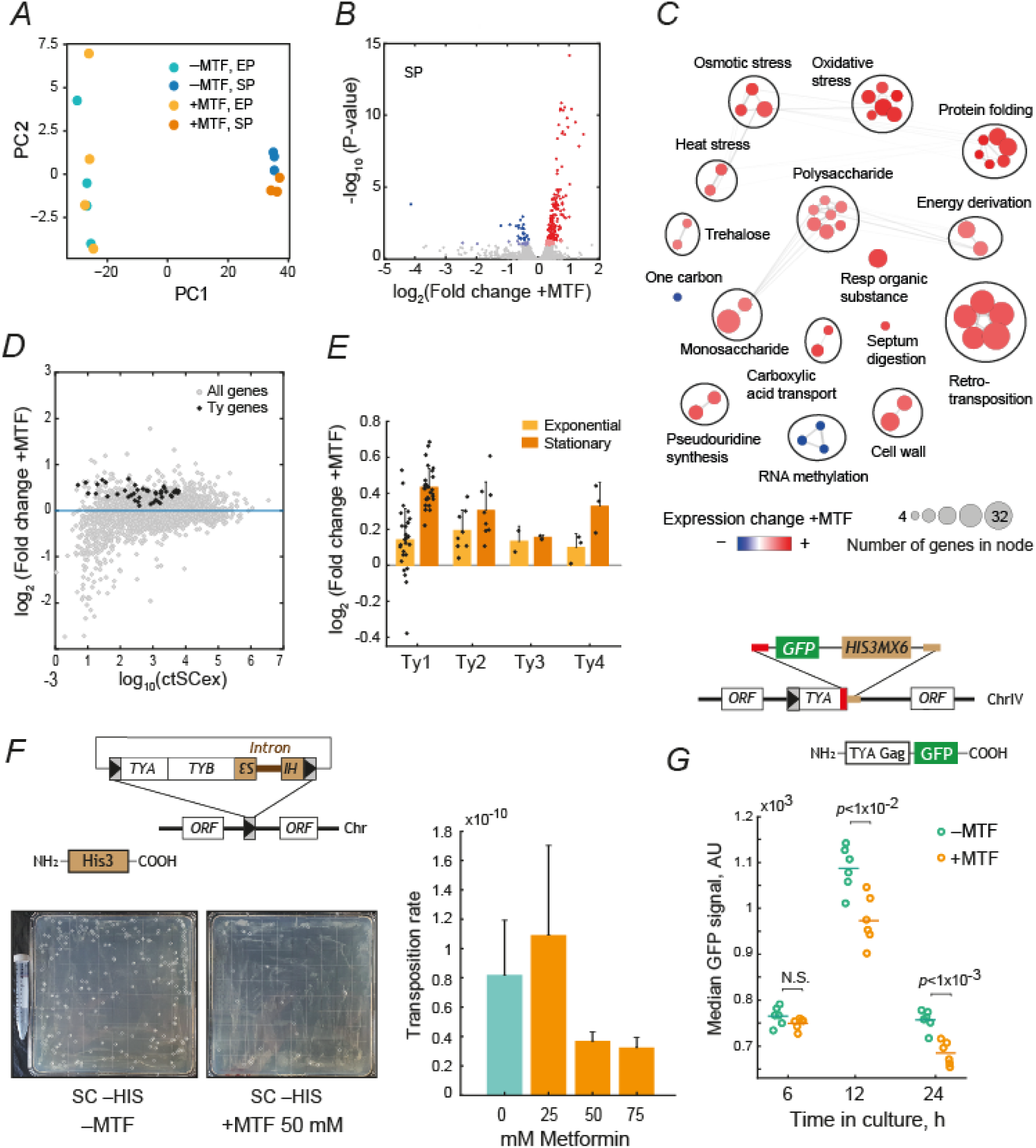
Changes in retrotransposon dynamics in response to metformin. ***A)*** Principal component analysis (PCA) of gene expression data of exponential (EP, *n*=3) and stationary phase groups (SP, *n*=4) without metformin (‒MTF) and treated with 50 mM metformin (+MTF). ***B)*** Volcano plot showing the differential expresses genes in response to 50 mM of metformin treatment (+MTF) in late stationary phase cells. Colored circles correspond to upregulated (red), or downregulated (blue) genes detected by DESeq2 analysis using a strict (*p*<0.05, dark dots) or lenient cutoff (*p*<0.1, light dots) after FDR correction. ***C)*** Functional clustering of upregulated (red) and downregulated (blue) genes group by mega-clusters. An enrichment analysis was conducted with differentially expressed genes. The network results from the functional clustering of biological processes found by similarity between groups. Each cluster is delimited by a circle and contains related gene groups. Node size represents the number of genes in the group. The width edge is constructed with the similarity Jaccard index estimated between groups. ***D)*** The scatter plot compares the expression of genes (gray) in response to metformin with its basal expression (expression in ‒MTF medium on exponential phase). Black points correspond to the Ty1 retrotransposon genes detected. ***E)*** Fold change of Ty retrotransposon families in exponential and stationary phase. ***F)*** Ty1 retromobility was measured employing a Ty1*his-AI* reporter strain treated with 0, 25, and 50 mM of metformin (*n*=14 for 0 and 50 mM and *n*=7 for 25 mM; *t*-test). **G)** The median intensity fluorescence of a Gag-GFP fusion was tracked at different time points (6, 12, and 24 h) in response to 0 or 50 mM of metformin (*n*= 6; *t*-test).

To provide a functional description of the biological processes involved in transcriptional reprogramming promoted by metformin, we conducted a GO enrichment analysis of the identified set of up and down-regulated genes (**Table S4**). GO groups were clustered based on their gene-content similarity, using the Jaccard index among the ensuing terms (**Figure 4C**). Our analysis showed that metformin resulted in increased expression of several gene sets with roles on stress response, such as osmotic stress, oxidative stress, heat shock and protein folding, carbohydrate metabolism, and trehalose accumulation. Strikingly, our results showed that the transcription of Ty retrotransposon genes increased when cells were aged in the presence of metformin (**Figure 4D**). A closer inspection of the transcriptional changes of different retrotransposon families revealed that this effect was stronger for members of the Ty1-copia group of retrotransposons (**Figure 4E**). Because Set3C components regulate Ty1 activity in yeast (Mou et al., 2006), our findings suggest that both impaired Set3C function and metformin treatment influence Ty1 expression, which may underlie the previously noted phenocopying of metformin-induced longevity by Set3C deletions.

### 3.5. Metformin result in decreased levels of retromobility and TYA Gag-like protein, despite increases in Ty1 transcriptional expression

Based on the observed increases of Ty1 retrotransposons expression upon treatment with metformin, we asked whether metformin also resulted in increased retromobility and integration into the genome. To answer this question, we used a plasmid-borne Ty1his3AI reporter bearing the sequence of the *YMRCTy1-3* element of *S. cerevisiae* (Hénault et al., 2020). After incubating WT cells at a permissive temperature for retrotransposition, we found that the transposition rate showed a modest two-fold decrease in cells treated with higher doses of metformin (**Figure 4F**). While this decreased rate of retromobility was not significant because of the high variability observed in untreated cells, it was surprising given that the same drug dose lead to increased Ty1 transcriptional expression.

To confirm the paradoxical pattern of robust Ty1 induction alongside a modest reduction in retromobility, we used the *YDR170-W* Ty1 locus as a TYA-GFP fusion reporter to track changes in the abundance of the TYA Gag-like protein in the transition to stationary phase. The TYA protein is analogous to the Gag protein of retroviruses and is essential for the formation of virus-like particles during the yeast retrotransposon replication cycle (Brookman et al., 1995). In agreement with reduced levels of retromobility, we found that treatment with metformin resulted in reduced levels of the TYA-GFP reporter, which was significant during post diauxic growth and early stationary phase at 12 and 24 hours in culture, respectively (**Figure 4G**). These results suggest that increased expression of Ty1 elements by metformin leads to increased genome stability defined as reduced rates of Gag-like protein and retromobility. Together, these findings reveal a previously unappreciated role for metformin in modulating retrotransposon activity, suggesting that enhanced Ty1 transcription may paradoxically contribute to genome stability by suppressing later steps of the retrotransposition cycle.

### 3.6. Metformin increases the abundance of proteins involved in mitochondrial function and stress-response

To further explore the cellular pathways through which metformin promotes longevity, beyond the genetic associations with Set3C and the transcriptional activation of Ty1 elements, we examined the dynamics of the proteome during the transition into stationary phase. This approach was aimed at identifying broad cellular targets of metformin that could contribute to downstream effects on retrotransposon activity and longevity. To this end, we used the GFP fusion collection coupled with flow cytometry to track changes in protein abundances (DeLuna et al., 2010; Newman et al., 2006). We first measured a subset of GFP fusions harvested at different times culture with or without metformin (**Figure S5**). The correlation between different replicate experiments was high at early stationary phase (24 and 48 h incubation) and decreased later in stationary phase (5 d), likely due to reduced GFP fluorescence signal during starvation (Roberts et al., 2016). Based on these observations, we screened the GFP collection focusing on cells at early stationary phase (24 h). Strains with undetectable fluorescence intensity in nominal conditions (Newman et al., 2006) were filtered out, resulting in a final dataset consisting of 2616 GFP fusion strains (**Table S5**). Finally, we estimated the median intensity fluorescence of each strain for the two conditions after removing doublets, which were the absolute protein expression levels of each gene in cells with and without metformin.

Our proteome analysis revealed that most GFP fusions were unaffected by metformin (**Figure 5A**). However, a subset of strains showed significantly increased or decreased reporter levels, such as the stress-induced cell wall glycoprotein Sed1 and the Gus1 Glutamyl-tRNA synthetase, respectively. We grouped the GFP fusion strains by their annotated GO term and compared the fold-change distribution of each group with the distribution of the 2616 strains tested. We found that metformin led to increased abundance of proteins with roles on removal of superoxide radicals, protein folding, and carbohydrate storage (**Figure 5B**), which agreed with our transcriptome screening. Likewise, proteins related to mitochondrial respiration and amino acid and fatty acid metabolism were also increased. Given that mitochondrial activity and ROS are known to impact Ty1 dynamics (Stoycheva et al., 2007; VanHoute and Maxwell, 2014), these changes may provide a link between metformin-induced mitochondrial remodeling and the observed upregulation of Ty1 elements. Conversely, the levels of proteins associated with DNA replication, RNA processing, filamentous-growth signaling, and aminoacyl-tRNA metabolism were reduced in response to metformin. The absence of clear functional groups related to Set3C or other chromatin-modifying complexes in our proteome dataset further supports the view that Set3C and metformin act indirectly through convergent effects, possibly mediated by changes in Ty1 expression. Overall, our proteomic survey indicates that metformin directly impacts mitochondrial activity, metabolism, and stress-response signatures, which emerge as early outcomes of metformin exposure.

**Figure 5.**
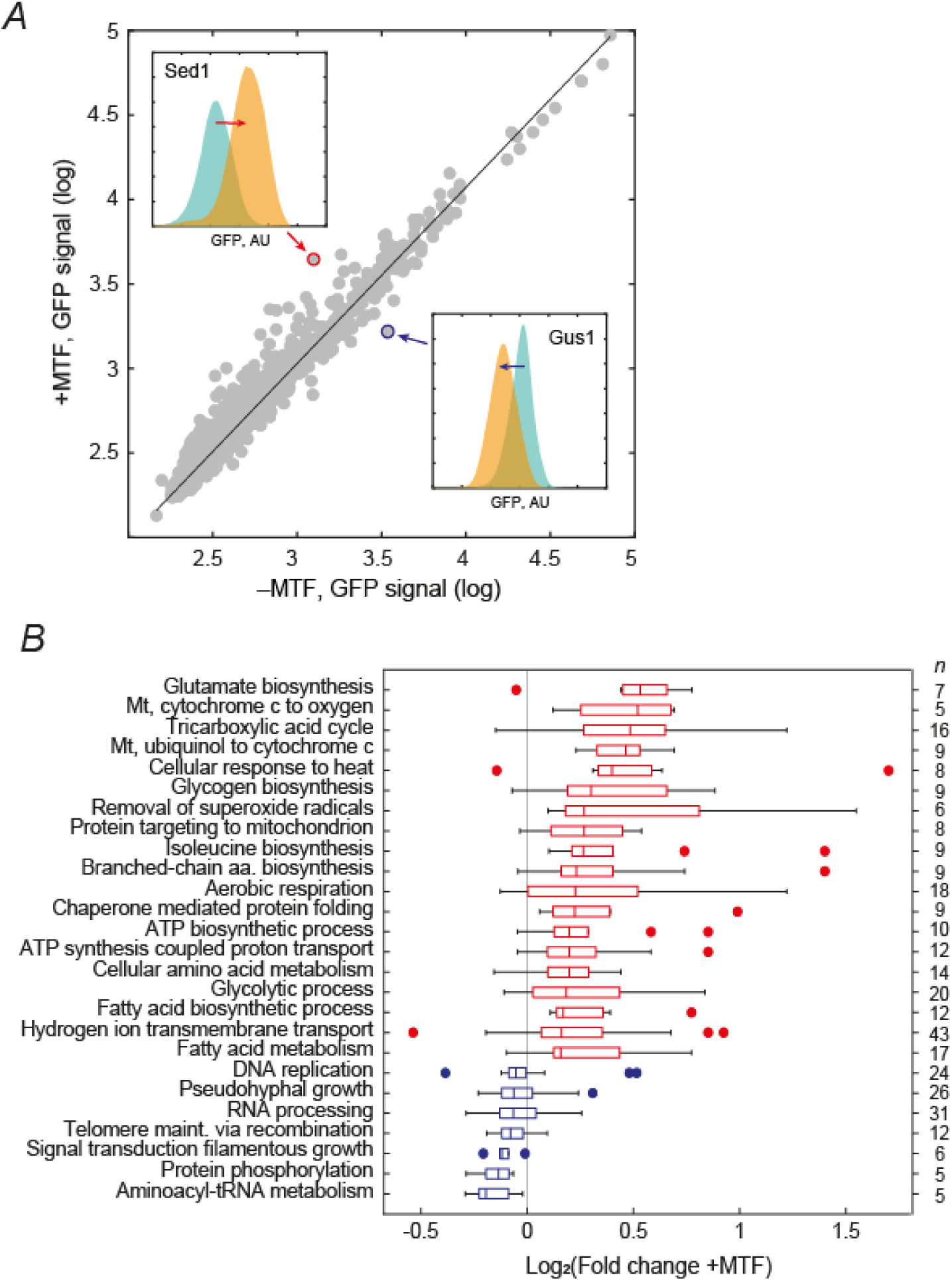
Proteomic response to metformin. ***A)*** Scatter plot of the GFP signal of 2616 fusion strains under no metformin (‒MTF, horizontal axis) and 50 mM metformin (+MTF, vertical axis), measured by flow cytometry. Insets show the GFP signal distribution of Sed1 and Gus1 in a media supplemented without metformin (cyan) and with 50 mM metformin (orange). ***B)*** Box plots show the fold-change distribution of the functional groups that change their abundance in response to metformin treatment (*p*<0.05, Wilcoxon rank sum test).

## 4. Discussion

Metformin is widely regarded as a promising compound with geroprotective potential (Barzilai et al., 2016; Kulkarni et al., 2020) with a strong safety profile and therapeutic benefits beyond its role as a first-line antidiabetic drug (Bu et al., 2022; Hua et al., 2023; Kasznicki et al., 2014; Lashen, 2010; Rotermund et al., 2018). Yet, despite knowledge of individual metformin targets, the pathways by which they converge to drive pro-longevity effects are not well understood. Here, we provide a comprehensive genetic and cellular analysis of metformin-induced lifespan extension in *S. cerevisiae*. Unexpectedly, we uncovered consistent gene–drug interactions between metformin and chromatin-modifying factors, including several components of the Set3C histone deacetylation complex. These results suggest that chromatin regulation is closely associated with metformin’s action on longevity. Notably, deletion of Set3C components extended lifespan in untreated yeast cells but diminished the pro-longevity effect of metformin, indicating convergence on shared pathways. To our knowledge, this is the first study linking metformin’s lifespan-extending effects to the Set3C histone deacetylation complex.

Aging is associated with heterochromatin loss and transcriptional activation of retrotransposable elements, leading to increased expression and mobility in multiple organisms, including humans (De Cecco et al., 2013; Hu et al., 2014; Li et al., 2013; Maxwell et al., 2011; Pal and Tyler, 2025; Patterson et al., 2015; Wang et al., 2011). It has also been hypothesized that metformin could act as a beneficial modulator of retrotransposon activity during embryological development (Finley, 2018). Our transcriptomic data showed elevated retrotransposon RNA levels in stationary-phase yeast cells; metformin further amplified this response, driving even higher transcription of Ty1 elements. Nevertheless, their retromobility was moderately reduced, suggesting a possible uncoupling of retrotransposon expression from their genome insertion. This mirrors observations in Set3C gene-deletion mutants, which also exhibit increased Ty1 RNA but reduced integration events (Mou et al., 2006), pointing to retrotransposon regulation as a shared outcome of Set3C impairment and metformin treatment.

The extent to which altered retrotransposon dynamics contribute causally to metformin-induced longevity remains unclear. Nonetheless, Ty1 retrotransposition has been previously associated with genome instability during chronological aging in yeasts (Maxwell et al., 2011). Further supporting a potential causal role, ectopic expression of Ty1 extends CLS in a wild *S. paradoxus* strain that naturally lacks retrotransposons, suggesting that retrotransposition can actively and directly promote longevity (VanHoute and Maxwell, 2014). Our finding that metformin increases Ty1 expression with minor effects in retromobility is consistent with recent studies in *Drosophila* showing that changes in retrotransposon expression can affect aging independently of genomic integration (Schneider et al., 2023). It also remains to be determined how metformin leads to increased Ty1 expression. Notably, metformin increased the abundance of proteins involved in mitochondrial function and in responses to ROS, both of which are known to modulate Ty1 dynamics in yeast (Stoycheva et al., 2007; VanHoute and Maxwell, 2014). Moreover, the Ty1-transcriptional activator Tec1 and mitochondrial function compensate for each other’s loss in determining chronological aging (Cruz-Bonilla et al., 2025), further supporting a link between metformin, stress responses, Ty1 regulation, and longevity in yeast.

By integrating large-scale genetic screening with proteomic and transcriptomic profiling, we uncovered both canonical responses to metformin and an unexpected link between lifespan extension, chromatin dynamics, and retrotransposon regulation. These findings expand the view of metformin’s cellular impact and its pro-longevity effects beyond metabolism and signaling pathways. As retrotransposons are increasingly recognized for their roles in genome regulation and stress adaptation, our study opens new avenues for exploring how metformin influences these elements and how such regulation contributes to aging and longevity across species.

## 5. Authorship contribution statement

J.M-P. and A.D. conceptualized and designed the study. J.M-P, E.M-M., C.P-A. and M.M-F. performed experiments. J.M-P, D.A. and A.D. performed data analysis. S.F. and A.D. oversaw experiments. C.A-G. and A.D. oversaw data analysis. S.F. and A.D. acquired funding. J.M-P. and A.D. wrote the original draft. All authors read and critically revised the manuscript.

## 8. Acknowledgements

We thank J. A. Avelar-Rivas, S. E. Campos, and E. Mancera for critical reading of the manuscript and Y. Macotela and A. Moreno-Estrada for helpful discussions. The yeast Ty1-retromobility reporter was kindly shared by C. R. Landry’s lab.

## 9. Data availability

All processed data are provided in the Supplementary Material accompanying this article. Raw RNAseq and flow cytometry data will be made available on NCBI GEO upon publication. The scripts used for data processing and analysis will be made available upon publication at https://github.com/.

## Abbreviations

C.F.U.: Colony-forming units
CFP: Cerulean fluorescent protein
CLS: Chronological lifespan
EDTA: Ethylenediaminetetraacetic acid
FDR: False Discovery Rate
GFP: Green fluorescent protein
GO: Gene ontology
*LE*: Lifespan extension coefficient
LTR: Long Terminal Repeat
MTF: Metformin
OD: Optical density
RFP: Red fluorescent protein (mCherry)
*S*: Relative survival coefficient
S.E.M.: Standard error of the mean
SC: Synthetic complete
Set3C: Set3 complex
SSC: Side scatter
WT: Wild type
YNB: Yeast nitrogen base
YPD: Yeast-Extract Peptone Dextrose

## Apendix A. Supplementary Material

Metformin-induced longevity is associated with retrotransposon dynamics in yest chronological aging.

### Supplementary Tables

**Table S1. Competitive aging output data and statistics (XLSX FILE)**

**Table S2. CLS-specific gene groups associated with metformin-induced longevity (XLSX FILE)**

**Table S3. RNASeq transcriptomic output data and statistics (XLSX FILE)**

**Table S4. GO enrichment analysis, RNAseq transcriptomics (XLSX FILE)**

**Table S5. GFP-fusions proteomic output data (XLSX FILE)**

### Supplementary Figures

**Supplementary Figure S1.**
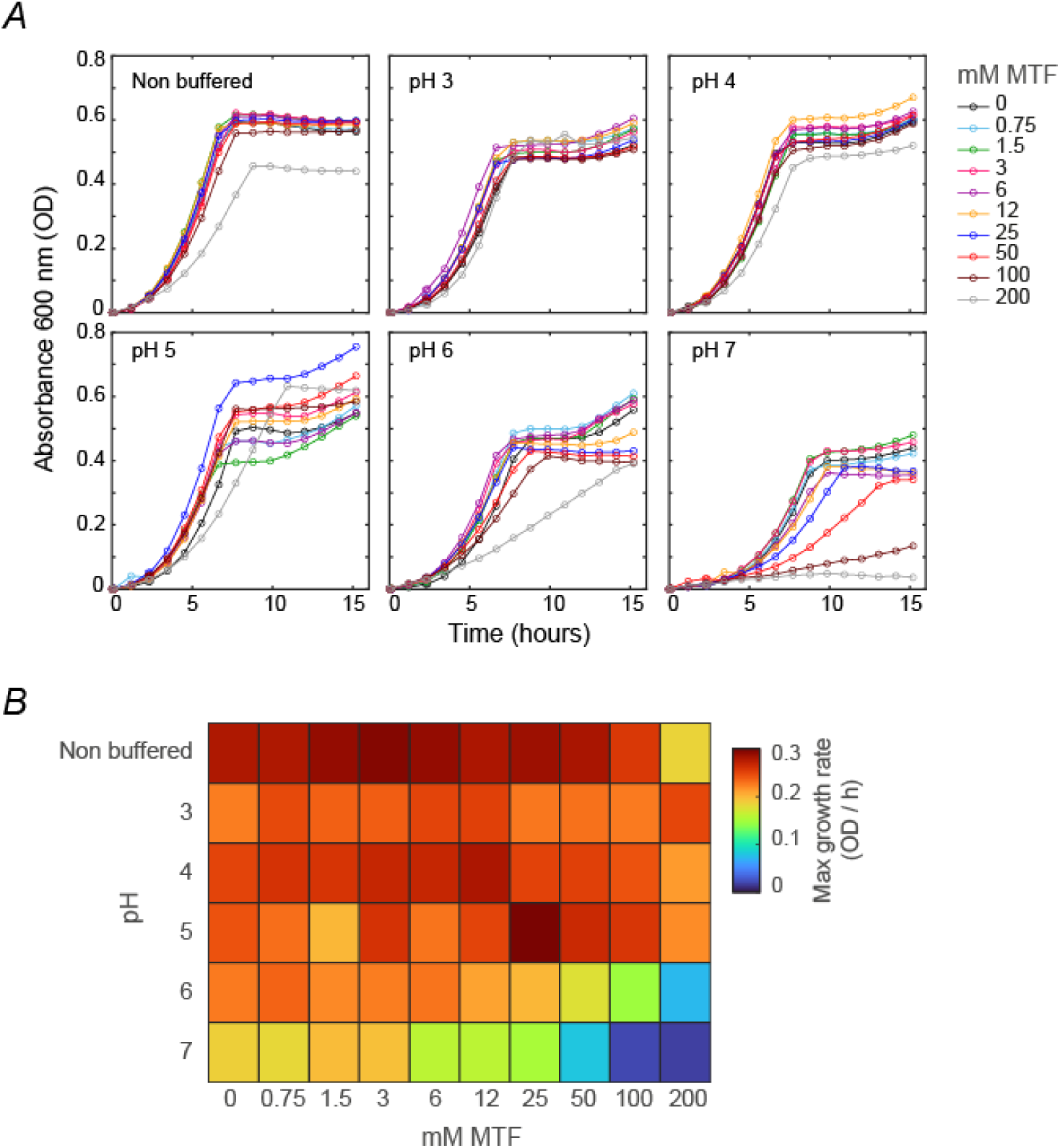
***A*)** Plots show growth kinetics of the WT strain at increasing concentrations of metformin, assayed from culture growth in non-buffered and buffered media adjusted at different pHs. ***B*)** Heatmap shows the maximum growth rates observed at increasing concentrations of metformin (horizontal axis) under different pH conditions (vertical axis).

**Supplementary Figure S2.**
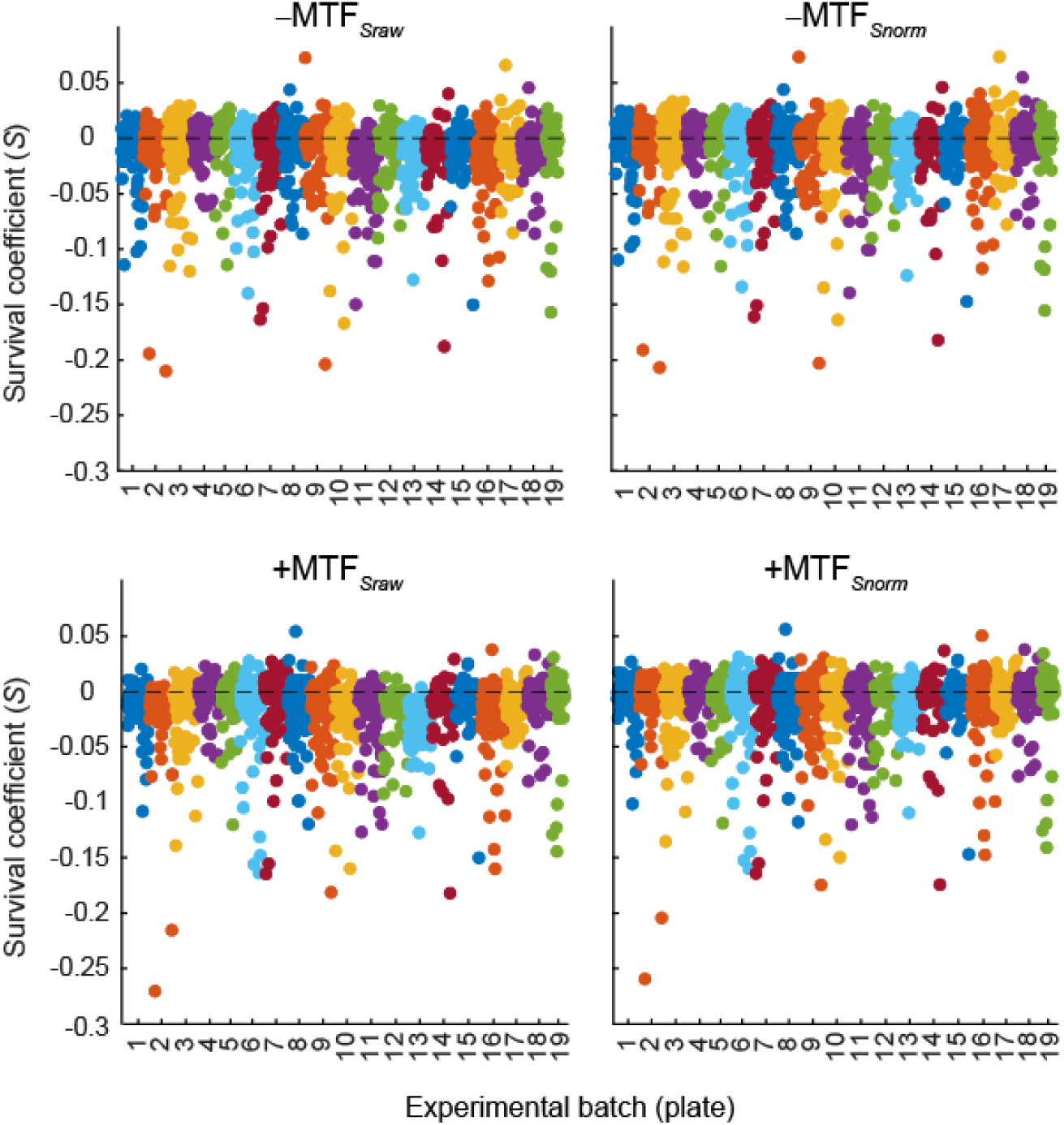
Raw (left panels) and normalized (right panels) survival coefficients (*S*) are plotted per plate for 0 mM (‒MTF) and 50 mM MTF (+MTF). To correct batch effects, data was normalized by subtracting the median *S* of each plate from the *S* of the mutants of the same plate, including WT reference replicates.

**Supplementary Figure S3.**
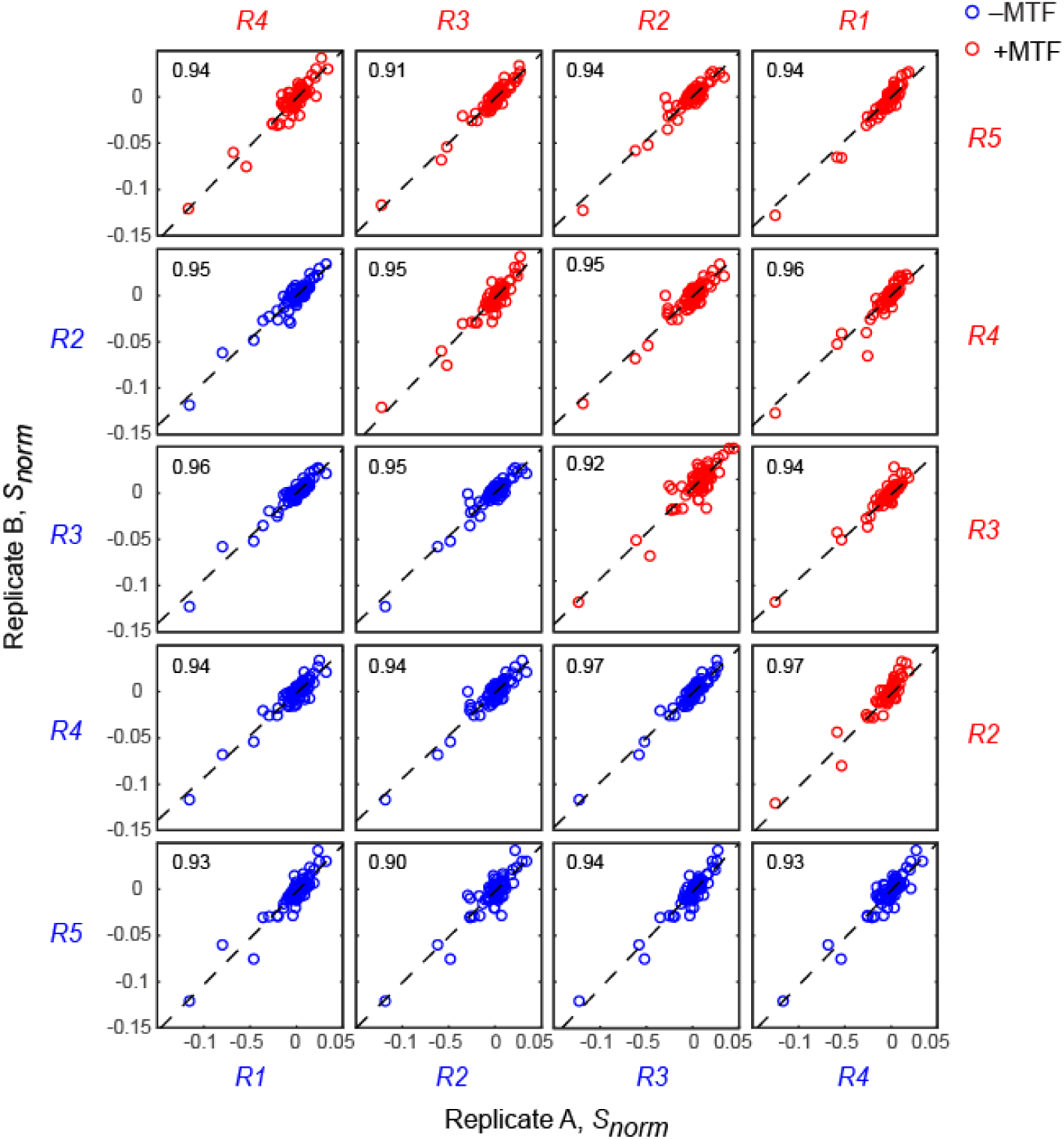
Scatterplot matrices of the survival coefficients of 84 mutants replicated in different batches to assess the replicability of our phenotyping approach. Each scatter plot also shows the correlation coefficient estimated for every replicate comparison (Pearson). Blue dots correspond to the 0 mM treatment (‒MTF) and red dots to 50 mM MTF (+MTF).

**Supplementary Figure S4.**
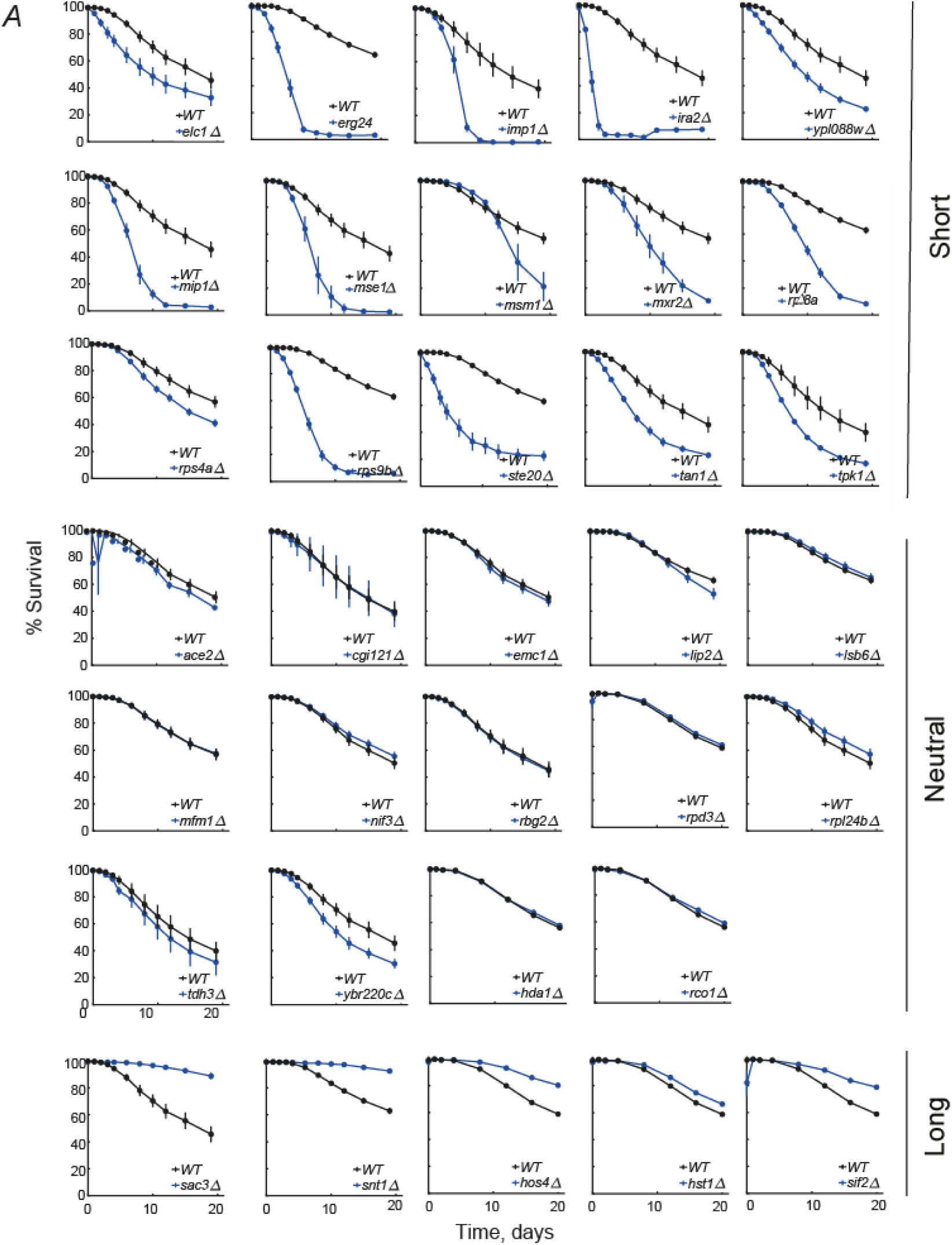
***A*)** Validation of CLS phenotypes in ‒MTF. Survival curves of selected mutant strains (blue lines, *n* = 5) were compared to WT (black lines) by live and death assay coupled to analytical cytometry. Error bars indicate the S.E.M. Upper and lower panels show short- and long-lived hits from the genetic screening, while middle panels show non-significant (neutral) phenotypes.

**Supplementary Figure S4.**
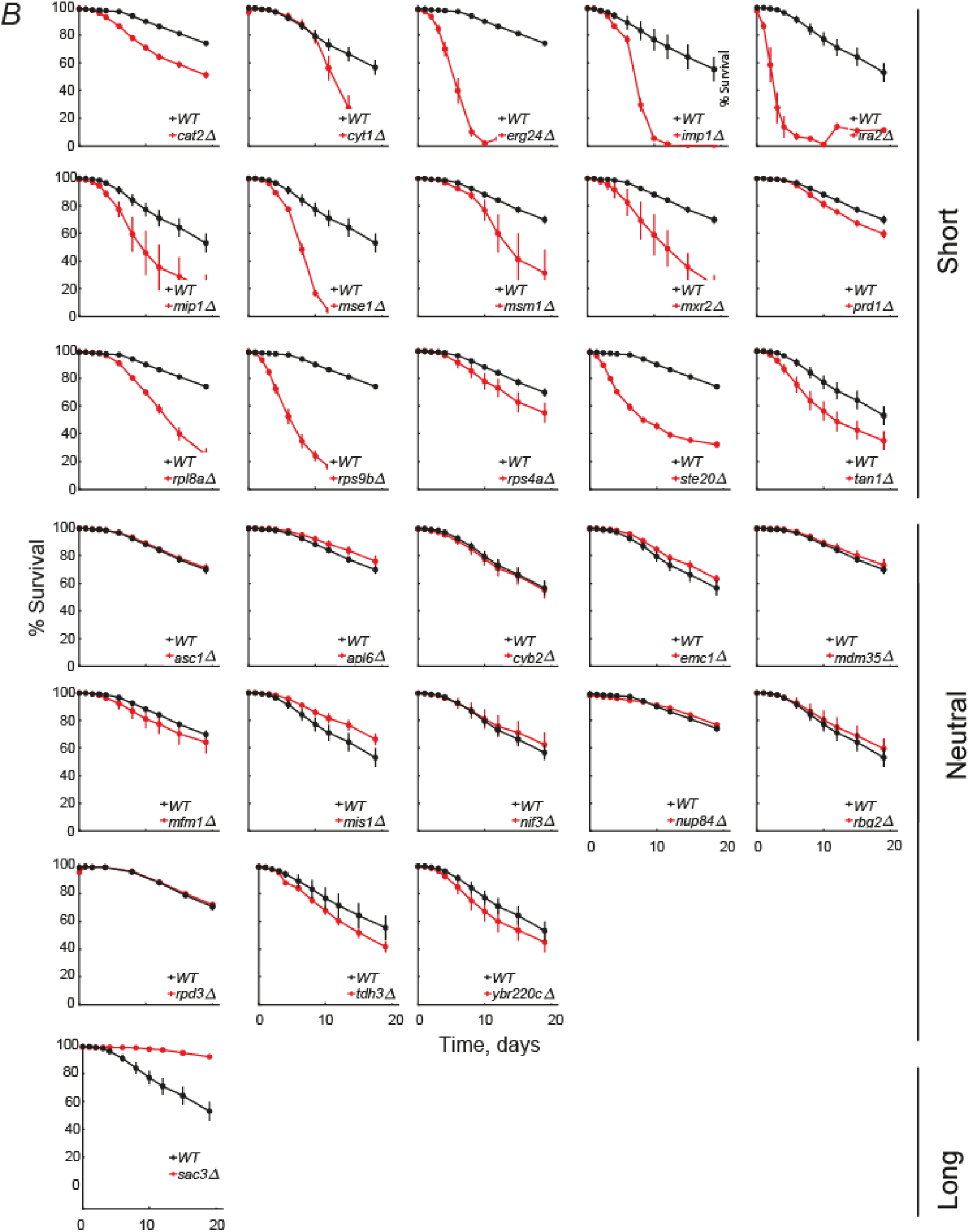
***B*)** Validation of CLS phenotypes in +MTF. Survival curves of selected mutant strains (red lines, *n* = 5) were compared to WT (black lines) by live and death assay coupled to analytical cytometry. Error bars indicate the S.E.M. Upper and lower panels show short- and long-lived hits from the genetic screening, while middle panels show non-significant (neutral) phenotypes.

**Supplementary Figure S5.**
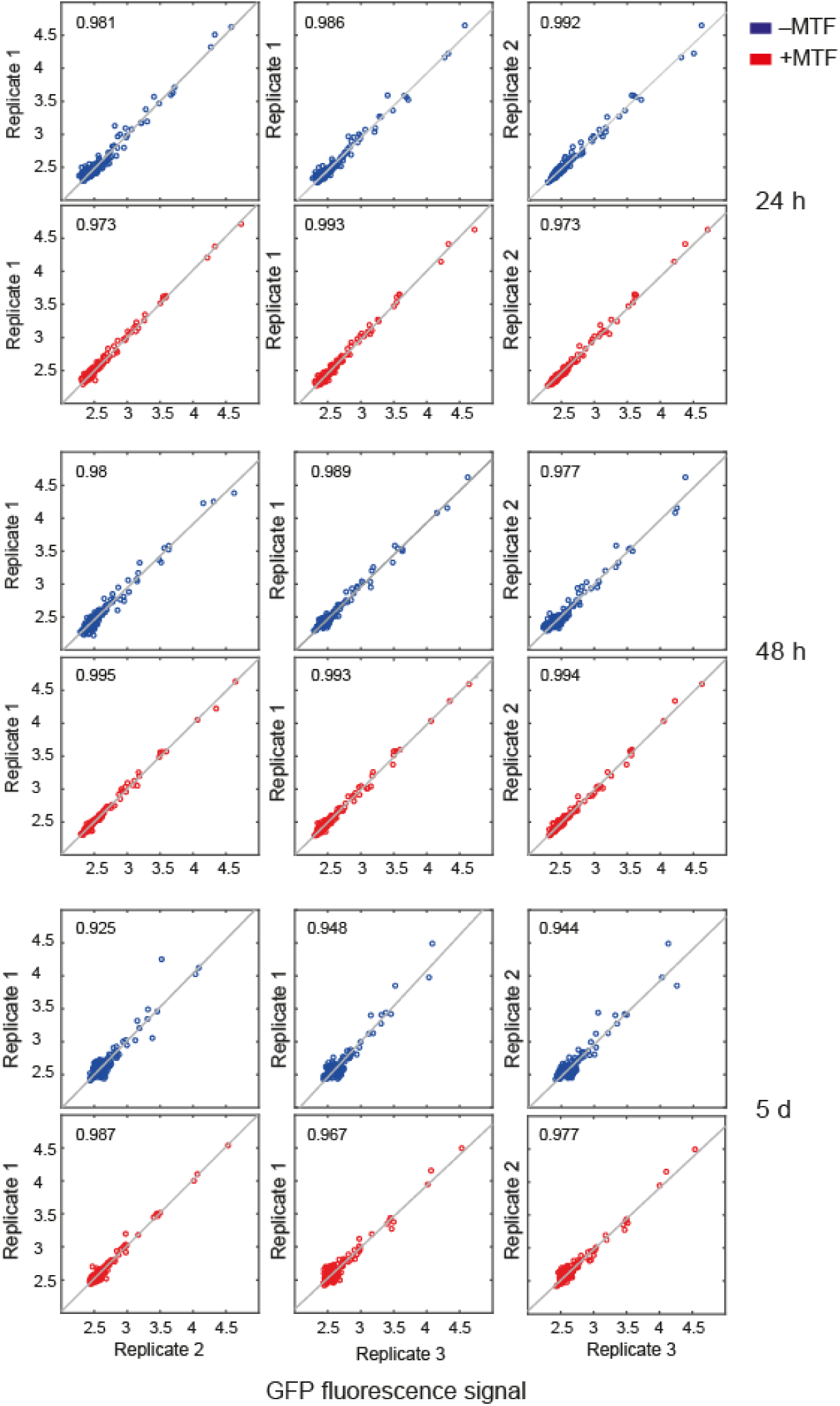
Scatterplot matrices of the median intensity fluorescence of 2616 GFP fusion strains replicated in different batches to assess the replicability of our method through time (24h, 48h and 5 days). Each scatter plot also shows the correlation coefficient estimated for every replicate comparison (Pearson). Expression levels in the no metformin (‒MTF) and 50 mM metformin (+MTF) treatments are shown in blue and red circles, respectively.

